# Volumetric Imaging of Human Mesenchymal Stem Cells (hMSCs) for Non-Destructive Quantification of 3D Cell Culture Growth

**DOI:** 10.1101/2022.09.19.508168

**Authors:** Oscar R. Benavides, Holly C. Gibbs, Berkley P. White, Roland Kaunas, Carl A. Gregory, Kristen C. Maitland

## Abstract

The adoption of cell-based therapies into the clinic will require tremendous large-scale expansion to satisfy future demand, and bioreactor-microcarrier cultures are best suited to meet this challenge. The use of spherical microcarriers, however, precludes in-process visualization and monitoring of cell number, morphology, and culture health. The development of novel expansion methods also motivates the advancement of analytical methods used to characterize these microcarrier cultures. A robust optical imaging and image-analysis assay to non-destructively quantify cell number and cell volume was developed. This method preserves 3D cell morphology and does not require membrane lysing, cellular detachment, or exogenous labeling. Complex cellular networks formed in microcarrier aggregates were imaged and analyzed *in toto*. Direct cell enumeration of large aggregates was performed *in toto* for the first time. This assay was successfully applied to monitor cellular growth of mesenchymal stem cells attached to spherical hydrogel microcarriers over time. Elastic scattering and fluorescence lightsheet microscopy were used to quantify cell volume and cell number at varying spatial scales. The presented study motivates the development of on-line optical imaging and image analysis systems for robust, automated, and non-destructive monitoring of bioreactor-microcarrier cell cultures.

## Introduction

In 2002, the Food and Drug Administration (FDA) announced a new science-based initiative to modernize quality management of pharmaceutical manufacturing and product quality by developing and implementing technologies that measure, control, and/or predict quality and performance of a process or product [1]. Utilizing Quality by Design (QbD) principles, traditional pharmaceutical and cell-based therapy manufacturers have recently been encouraged to develop and utilize Process Analytical Technologies (PATs) that perform real time or near real time monitoring of key variables throughout the manufacturing process to ensure quality of the final pharmaceutical product [2], [3]. In cell-based therapy manufacturing, PATs are designed to increase understanding of the cell culture process and facilitate the monitoring and control of critical process parameters (CPPs) that directly influence the quality and safety of the final cell product [4]. Ideal systems perform automated, (near-) real time, on- or in-line, non-destructive measurements and analysis. The identification of CPPs for stem cell cultures and the development and deployment of PATs will improve the ability to study novel cell lines and expansion methods and enhance the capability to monitor, control, and ultimately predict the final product quality [2], [5].

Cell-based therapies, or cytotherapies, have the potential to address an unmet need for therapies that cure or treat chronic diseases such as cancer, osteoporosis, diabetes, and stroke [6]–[9]. Entry into the clinic will require billions of cells per indication per year, and one critical challenge in upstream cytotherapy manufacturing is the efficient large-scale expansion of stem cells to maximize yield while maintaining safety and therapeutic efficacy [10]. Even with the latest generation of multi-stacked cell factories, two-dimensional (2D) monolayer cultures have limited surface area for expansion, are labor- and reagent-intensive, and require serial passaging, which renders them sub-optimal for large-scale cellular expansion [11]. Three-dimensional (3D) cell cultures better mimic the *in vivo* stem cell niche than standard monolayer cultures while exploiting the 3^rd^ spatial dimension for cellular expansion [12]–[14]. Bioreactor-microcarrier suspension cultures are the most promising 3D cell culture method as bioreactors can be scaled up to volumes of 1,000 liters, and microspheres greatly increase the available surface area to volume ratio, reduce labor and reagent use, and can be functionalized for specific needs [10], [15]–[18]. Furthermore, several groups have shown that 3D microcarrier human mesenchymal stromal cell (hMSC) cultures can provide a greater cell yield than 2D monolayer cultures without compromising viability, identity, or differentiation potential [19]–[23]. Recently, our group used spherical biodegradable gelatin methacryloyl (gelMA) microcarriers and bioreactor suspension cultures to demonstrate scalable expansion, rapid harvest, and non-destructive 3D *in toto* sub-micron visualization of induced pluripotent stem cell-derived hMSCs (ih-MSCs) via reflectance confocal microscopy (RCM) without the need for detachment from the microcarrier surface [24].

In regenerative medicine and biopharmaceutical manufacturing involving expansion and harvest of living cells, cell number is the most fundamental cell culture process parameter that requires quantification. Cell enumeration is needed to evaluate viability and proliferation, and in functional assays where activity is normalized to cell number such as engraftment [25], [26].While there is no single established cell enumeration method for microcarrier cultures, essentially all off-line methods are destructive as they require either detachment of cells from the microcarrier surface and/or membrane lysing and exogenous labeling [27]. Two of the most common off-line cell enumeration and viability assays, trypan blue dye exclusion and live/dead fluorescence using Calcein AM and propidium iodide (PI), both of which are based on membrane integrity, are destructive to the samples and remove potentially valuable morphological and spatial distribution information [28]–[34]. The chemical analysis of off-gas, media, protein, or DNA content in a sample can be an indirect measure of cell concentration performed off-line following bioreactor sampling or on-line with *in situ* sensors [35]–[39]. Additionally, cell number can be measured on-line or in-line via a number of optical techniques, such as optical density measurements, *in situ* microscopy, micro-flow imaging, imaging and flow cytometry, and IR and fluorescence spectroscopy, and non-optical methods based on dielectric spectroscopy or acoustic measurements [37], [40]–[53].

Non-visualization cell enumeration methods are incapable of providing insight on cell morphology or spatial distribution, which in traditional monolayer cultures are readily monitored using in-process brightfield or phase-contrast microscopy; these CPPs are informative and potentially predictive features of cellular fate, proliferation, and functional potential in monolayer cultures [4], [5], [30]–[34], [54], [55]. The evaluation of microcarrier surface confluency and spatial distribution of cells can provide insight into the culture microenvironment and better enable automated, objective real-time release of intermediate upstream cell cultures once a certain confluency threshold is reached [30], [56]. An ideal biomass monitoring PAT for bioreactor-microcarrier anchorage-dependent cell culture performs measurements on-line or inline and *in toto*, leaving cells attached to microcarriers and cell-microcarrier aggregates undisturbed so as to preserve cell morphology and 3D spatial distribution information.

Several imaging, microscopy, and visualization methods for cell enumeration in microcarrier-based cultures have been investigated, but an industry standard has yet to be determined. Automated image analysis could be incorporated to image- or visualization-based assays for more rapid and robust quantification [57]–[59].Trypan blue dye exclusion and live/dead fluorescence labeling both require exogenous contrast agents, so they cannot be incorporated into on- or in-line assays. Off-line fluorescence-based direct cell enumeration assays are, however, used to correlate experimental on-line cell enumeration or biomass sensors [35], [60]. These assays have been based on total fluorescence intensity as opposed to 3D spatial volume which considers 3D cell morphology. Volumetric fluorescence microscopy can be used to characterize cell density, distribution, and morphology, but requires destructive exogenous fluorescent markers [61], [62].We previously demonstrated RCM, based on back-scattered elastic photons, could be employed to achieve label-free, sub-micron *in toto* visualization of hMSCs attached to spherical microcarriers [24]. This optical method allows for cell enumeration, but raster scanning a 3D volume of ∼150^3^ μm^3^ is too slow for on-, in-, or even off-line measurements. The rapid and photo-efficient light sheet microscopy (LSM) technique presents a more viable method for non-destructive monitoring of microcarrier cultures [63], [64]. Fortunately, elastic scattering light sheet microscopy (esLSM), also known as light sheet tomography (LST), can be used for *in toto* imaging of hMSCs attached to microcarriers, quantification of cell number while preserving cell morphology. Contrast is generated from elastically scattered photons as opposed to more traditional light sheet fluorescence microscopy (LSFM) that utilizes fluorescence for imaging [65]–[67].

Here, we report a proof-of-concept study on volumetric optical imaging and semiautomated image analysis for off-line fluorescence and on-line elastic scattering quantification of cell number and volume of hMSCs cultured on spherical hydrogel microcarriers *in toto*, without the need for cellular detachment. The off-line fluorescence assay utilizes LSFM and CellTracker Green cytoplasmic and DRAQ-5 nuclear labeling for cell volume quantitation and direct cell enumeration of single microcarriers and large aggregates. With this fluorescence-based assay, cell number was correlated to cell volume estimated from the elastic scattering-based assay. The presented off-line assay is the first imaging-based assay to use volumetric data to more accurately characterize the 3D microcarrier cell culture and the first to directly enumerate cells within large aggregates *in toto*. We demonstrate that the esLSM and image analysis (ELIAS) method has the capability to be adopted as an on-line PAT for robust non-destructive monitoring of bioreactor-microcarrier cell culture growth, and would improve process and quality control for cytotherapy manufacturing.

## Material and Methods

### Induced pluripotent stem cell-derived hMSC (ih-MSC) culture

Passage 4 ih-MSCs were first expanded in low-density monolayer cell culture in complete culture medium (CCM) (α-Minimum Essential Medium, 10% fetal bovine serum, 2 mM L-glutamine, 100 U/mL penicillin, and 100 μg/mL streptomycin) to obtain the required cell numbers. The ih-MSCs [68] were cultured in a rotating wall vessel (RWV) bioreactor (RCCS-8DQ bioreactor (Synthecon, Houston, TX) fitted with 10 mL high aspect ratio vessels [69] on custom-fabricated 120 ± 6.2 μm diameter gelMA microcarriers [24]. For this purpose, approximately 110,000 gelMA microcarriers with a combined growth area of 50 cm^2^ and 5×10^4^ cells (1000 cells/cm^2^) were incubated in 10 mL of CCM in the RWV bioreactor at 24 revolutions per minute. Half of the media was replaced with fresh CCM every 2 days. Specimens were recovered and fixed at passages 4 and 7 on days 3 and 7, for a total of four samples.

### Sample preparation

At day 3 and day 7 of RWV bioreactor culture, CCM was removed and microcarrier-expanded cells were suspended in 1 mM concentration of CellTracker Green (CTG) for 45 minutes. The CTG target is distributed uniformly in the cell cytoplasm, and was used here to visualize 3D cell morphology and quantify cell volume. Cells were fixed with 4% paraformaldehyde (PFA) and stored in phosphate buffered saline (PBS) at a concentration of 3 mg particles/mL PBS for long-term storage. Fixed microcarrier-cell samples were incubated with a 5 μM DRAQ-5 DNA and 6.5 μM DiI plasma membrane staining buffer at 37º C for 30 minutes with agitation, then rinsed with PBS. The far-red fluorescent DRAQ-5 stain was used for cell nuclei visualization and direct cell enumeration. The orange fluorescent DiI label was used to illustrate a simpler staining method for visualization of the plasma membrane only.

The microcarriers were embedded in 1% agarose in a custom-designed and 3D-printed sample chamber (S1 Fig.) [70]. A 300 μL aliquot was loaded into each sample chamber at a concentration of 6 mg particles/mL agarose. The chamber enables dual-sided lightsheet illumination, trans-illumination for widefield imaging, and >180° sample rotation for multi-view acquisition and optimized sample positioning.

### Off-line fluorescence-based cell enumeration and cell volume quantification

The Zeiss Z1 Lightsheet microscope, with a 20X 1.0 NA (water) detection objective lens and 10X 0.2 NA illumination objective lenses, was used for *in toto* imaging of fixed ih-MSCs attached to spherical microcarriers. The 488 nm (power 5%) and 638 nm (power 9%) lasers were used to excite the CTG and DRAQ-5 fluorophores, respectively. The voxel size was 0.2 × 0.2 × 0.45 μm^3^ to satisfy Nyquist sampling requirements. The emission filters used were 505 – 545 nm and 660+ nm for CTG and DRAQ-5, respectively. Dual objective illumination with pivot scanning and online max fusion was used to improve illumination of microcarrier aggregates and reduce acquisition time. The camera integration time was set to 20 ms per frame, or 50 FPS, and the illumination power was adjusted to use the full dynamic range of the detector.

Imaris image analysis software was used to view and analyze the 3D volumes. For direct cell enumeration, the ‘Spot’ function was used on the DRAQ-5 volumes with an object size filter of 10 μm and high-pass intensity threshold to segment and enumerate cell nuclei (S2 Fig.). For quantification of cell volume, the ‘Surface’ function was used on the CTG volumes with a high-pass intensity threshold and size filter to remove cell debris from quantification (S3 Fig.). No preprocessing of the data was required for cell segmentation as the gelMA microcarriers produce little background signal [24]. For enumerating microcarriers, the ‘Surface’ function was used on the CTG volume with a low-pass intensity-based threshold to create a solid object. Then, the ‘Spot’ function with a 90 μm size filter was used to identify individual spherical microcarriers.

### On-line elastic scattering-based cell volume quantification

The agarose-embedded and mounted microcarrier-cell samples were used to evaluate the feasibility of the ELIAS method for label-free, non-destructive, *in toto* imaging and characterization of 3D microcarrier cell culture growth.

For elastic scattering imaging on the Z1 Lightsheet microscope, the 638 nm laser (power 0.1%) was used to illuminate the sample. The laser blocking filter and emission filters were removed from the optical path. The camera acquisition time was minimized to 10 ms per frame, and the laser power was adjusted until there were no saturated pixels when viewing a cell. Dual-sided illumination, pivot scanning, and online max fusion were turned on. The Imaris ‘Surface’ function was used to segment the cells and microcarriers from each other and the agarose. For cell volume quantification, a high-pass intensity threshold and a high-pass size filter were used (S4 Fig.). For microcarrier enumeration, a low-pass intensity threshold and 90 μm size filter were used (S5 Fig).

## Results

### 3D visualization via lightsheet microscopy

To demonstrate the ability to use volumetric fluorescence and label-free elastic scattering microscopy for direct and non-destructive quantitative monitoring of cell culture growth, ih-MSCs attached to gelMA microcarriers were imaged at four timepoints using lightsheet microscopy (Figure 1). The gelMA microcarriers, which have a refractive index (n) of 1.35 [24], permit 3D visualization of the entire microcarrier surface and core, enabling direct cell enumeration *in toto* via DRAQ-5-labeled nuclei and quantification of cell volume using LSFM (S1 and S2 Videos) and esLSM (S3 and S4 Videos) while preserving the integrity of the cell morphology.

**Figure 1.**
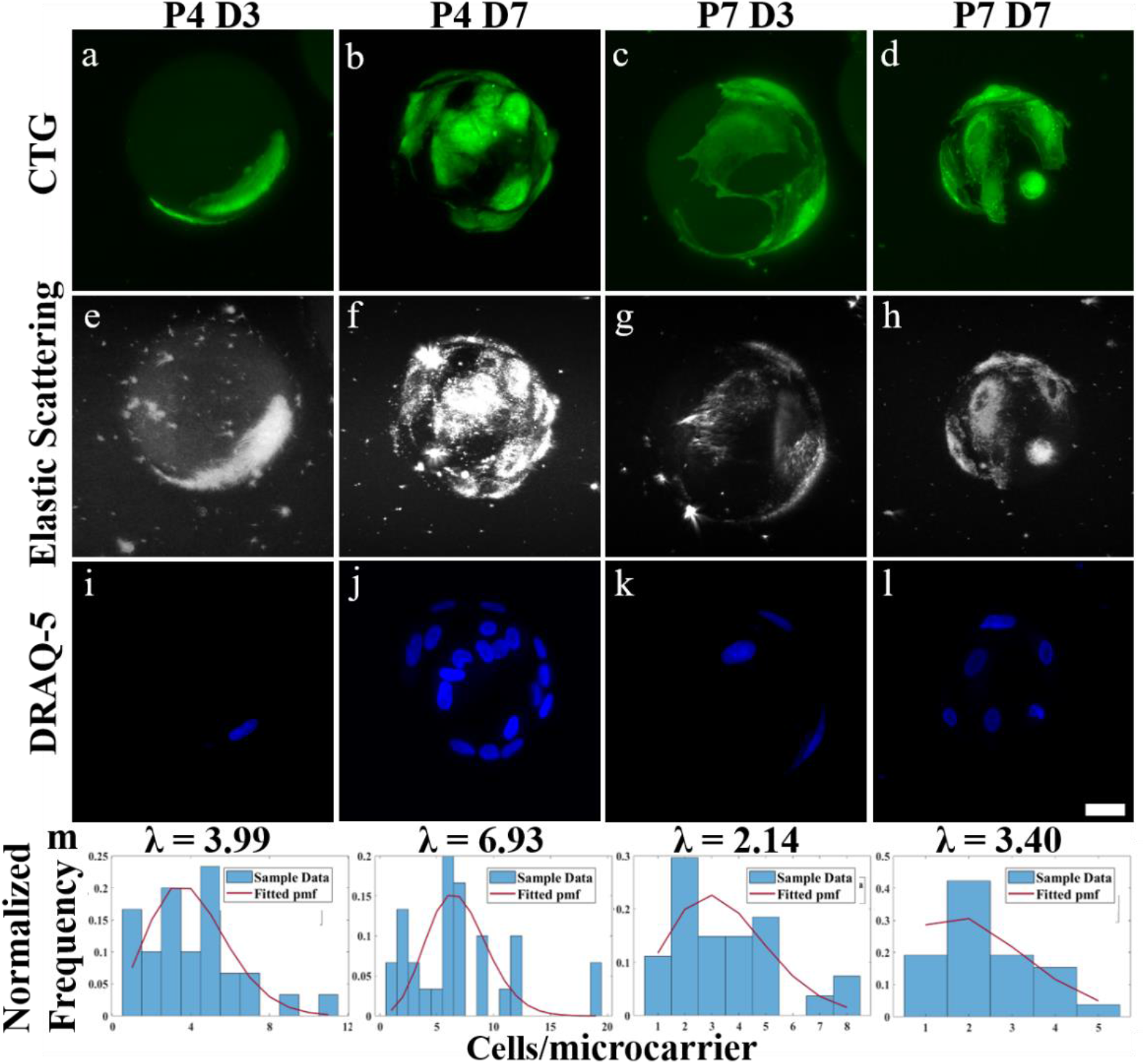
LSM and hydrogel microcarriers enable direct cell enumeration (DRAQ-5) and cell volume (CTG) quantification *in toto*. Representative 2D LSM max. intensity projections of ih-MSCs attached to gelMA microcarriers at passage 4 day 3 (P4 D3), passage 4 day 7 (P4 D7), passage 7 day 3 (P7 D3), and passage 7 day 7 (P7 D7) using **a-d)** CTG fluorescence and **e-h)** elastic scattering contrast. **i-l)** DRAQ-5 labeled nuclei were used to estimate the **m)** normalized frequency of counted ih-MSCs attached to single gelMA microcarriers at P4 D3 (n = 30), P4 D7 (n = 20), P7 D3 (n = 23), and P7 D7 (n = 26). A zero-truncated Poisson distribution is fit over the sampled data (red line). Scale bar = 25 μm.

The CTG volumes reveal the individual cell morphologies, and there appeared to be an increase in total cell volume from day 3 to day 7 in both passages (Figure 1a-d). The label-free elastic scattering shows similar cell morphology and cell growth over time as the fluorescence data (Figure 1e-lh). This suggests both contrast methods can be used to view cells and quantify cell volume. The elastic scattering signal seems to originate from the cytoplasm, and the nucleus tends to appear as a cavity that exhibits limited signal (Figure 1h). The DRAQ-5-labeled nuclei data show an increase in cell density from day 3 to day 7 within each passage (Figure 1i-l). The DRAQ-5 volumes were used to monitor the distribution of cells/microcarrier on single microcarriers over time (Figure 1m). Aggregates were excluded from the histograms to avoid weighting against single microcarriers. Similarly, non-populated microcarriers were not studied. A zero-truncated Poisson distribution was fit to the histograms to account for excluding empty microcarriers from acquisition and analysis [71]. These data suggest a decrease in cells/microcarrier or cell density at passage 7 compared to passage 4 overall. This cytoplasm-dominant elastic scattering phenomena has been previously reported [72], and here we similarly show that nuclear-bound fluorescent markers and elastic scattering microscopy provide complimentary information on different cell regions.

### Optical sectioning and hydrogel microcarriers

In LSM, a stack of thin (∼2 μm) planes is sequentially illuminated within the microcarrier sample, allowing more precise localization of interesting higher-resolution biological phenomena, such as a cell infiltrating the center of a gelMA microcarrier (Figure 2). The infiltration is clearly discernible in a 3D rendering of the dataset using both fluorescence and elastic scattering contrast (S5 and S6 Videos). In the 2D max. intensity projection of the microcarrier 3D volume from the LSM, it is not possible to discern the cell process burrowing into the core of the microcarrier (Figure 2a). This appears to be a large, binucleated cell wrapping around ∼1/3 of the microcarrier (Figure 2b). A max. intensity projection of the middle 1/3 volume of the microcarrier allows for visualization of the cell process extending into the core of the microcarrier (Figure 2c). There is, interestingly, a clear delineation of the nuclear envelope and microcarrier infiltration outlined by DiI staining (Figure 2d) [73]. Assuredly, the elastic scattering mode is also able to visualize this cell infiltration, illustrating that complex cell-microcarrier interactions and cell morphologies can be visualized and characterized with esLSM (Figure 2e).

**Figure 2.**
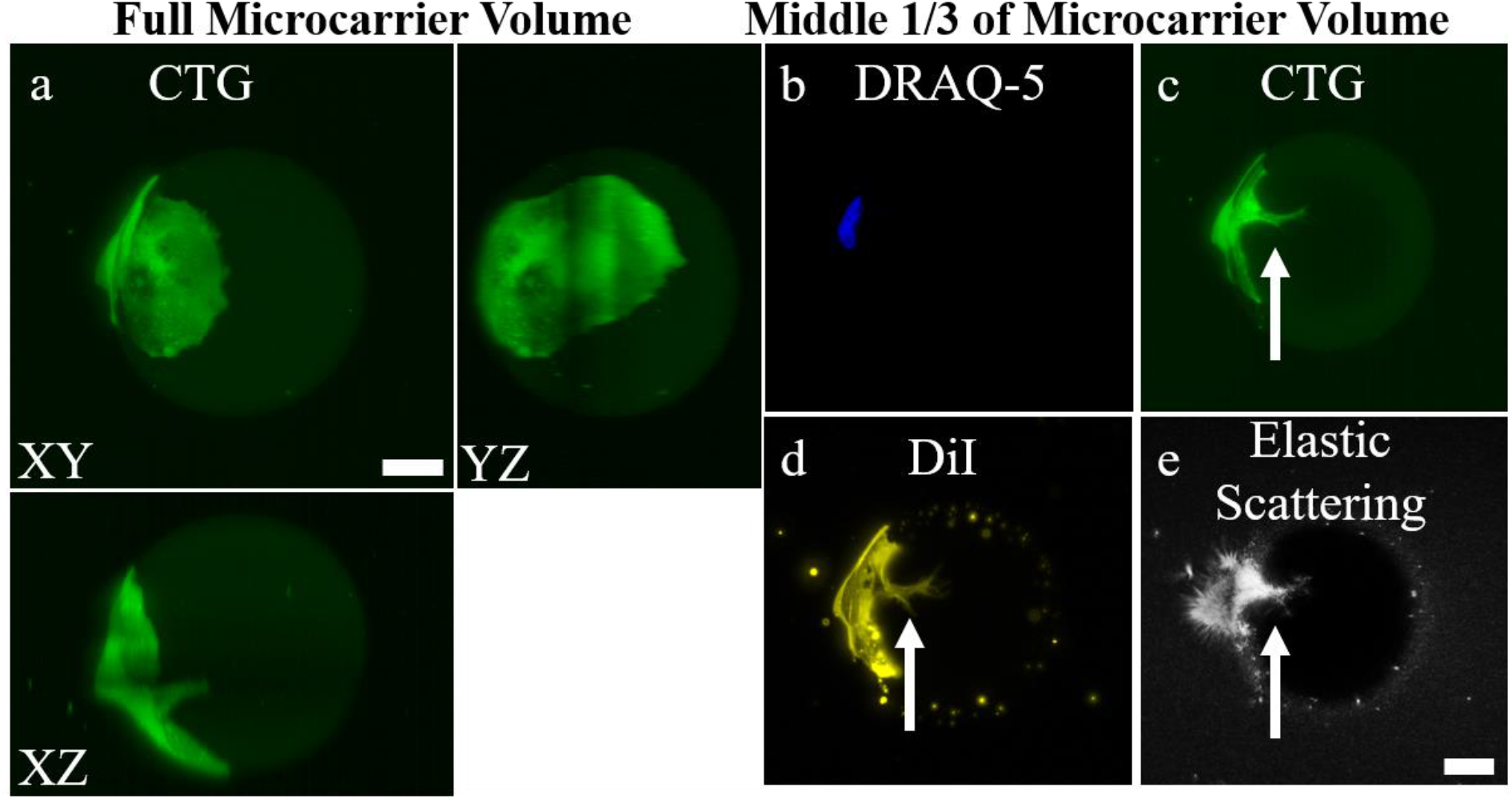
The optical sectioning capabilities of LSM permits visualization into the hydrogel microcarrier. **a)** 2D max. intensity and orthogonal projection of P7 D3 microcarrier with a single CTG-expressing cell. 2D max. intensity projections using the middle 1/3 of the volume to view the interior of the microcarrier using **b)** DRAQ-5, **c)** CTG, **d)** DiI plasma membrane stain, and **e)** elastic scattering contrast. Owing to the optical sectioning capabilities of LSM and refractive index matching of the microcarriers, biological features such as cell infiltration into the microcarrier can be visualized (white arrow). Scale bar = 25 μm.

### *In toto* imaging of large aggregates

Owing to the superior optical properties of the hydrogel microcarrier and optical sectioning ability of LSM, large cell-microcarrier aggregates that form later in culture with increased cell growth can reach volumes > 4 mm^3^ and still be imaged *in toto*. This permits semi-automatic cell enumeration, microcarrier enumeration, and cell volume quantification of large microcarrier aggregates using both off-line fluorescence and the ELIAS methods (Figure 3). The CTG data shows a complex network of cellular connections throughout the aggregate (Figure 3a). The DiI plasma membrane stain, which does not require a live incubation period for conversion into a fluorescent marker, similarly reveals a large cellular network (Figure 3b). There were 5,673 individual cell nuclei enumerated using the DRAQ-5 data (Figure 3c). Using the elastic scattering data, which provides slight contrast for the gelMA material, 1,754 microcarriers were detected, for an average of 3.32 cells per microcarrier (Figure 3d). The elastic scattering modality also reveals the cells throughout the entire microcarrier aggregate (Figure 3d). Small scatterers in the agarose and cell debris on the microcarrier surfaces can be segmented out with intensity- and size-based filters as cells scatter at higher intensity values and are larger than the debris (Figure 3e). The higher-resolution, merged projection of the DRAQ-5 and CTG data illustrates the density of cells within aggregates (Figure 3f). The elastic scattering and DRAQ-5 zoomed-in merged projection reveals similar cell morphologies as the CTG data even for this large aggregate (Figure 3g). A 1 mm sweep in depth through the aggregate further exemplifies that elastic scattering can visualize both microcarriers and the complex network of cellular connections that create an aggregate in culture (S7 Video).

**Figure 3.**
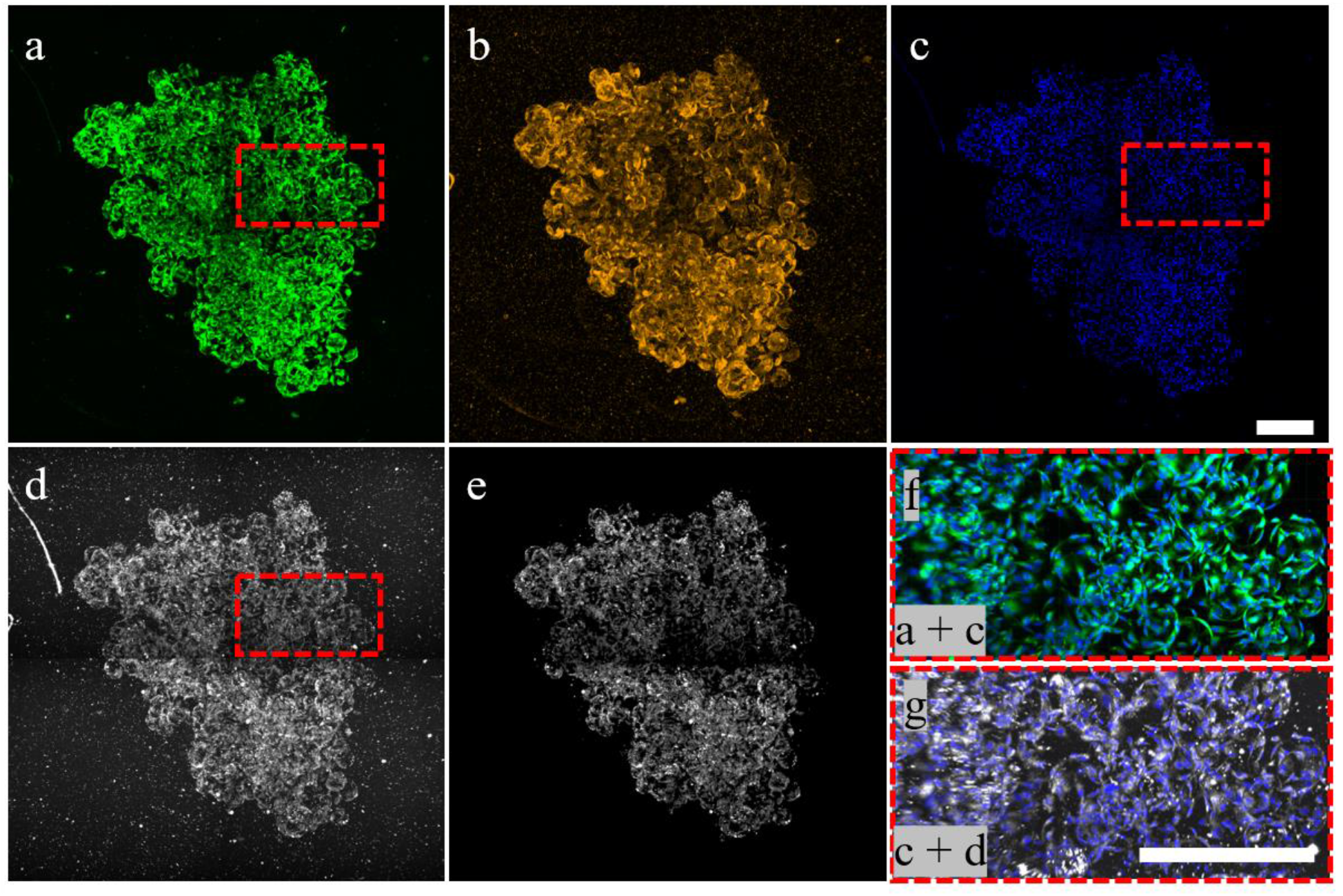
Large cell-microcarrier aggregates can be imaged and analyzed *in toto*, performing direct cell and microcarrier enumeration and quantifying cell volume using fluorescence and label-free LSM. Minimal refractive index mismatch between the hydrogel-based microcarriers and surrounding medium enable high resolution visualization of cell morphology throughout the entire aggregate. 2D maximum fluorescence intensity Z-projections of ih-MSCs labeled with **a)** CTG, **b)** DiI, and **c)** DRAQ-5. **(d)** Elastic scattering also allows for visualization of cells within large aggregates. **e)** Small scatterers in agarose and cell debris can be segmented out based on intensity and/or size using Imaris. **f)** CTG + DRAQ-5 merge and **g)** elastic scattering + DRAQ-5 merge at higher resolution. Scale bar = 400 μm.

### Cell enumeration and cell volume quantification

An off-line fluorescence-based assay for direct cell enumeration and cell volume quantification of cell expansion on microcarriers was developed. This method is based on the volume of cellular fluorescence as opposed to total fluorescence intensity. The DRAQ-5 data show that the average cells/microcarrier increased from day 3 to day 7 during both passages, but at a greater rate during passage 4 than passage 7 (Figure 4a). However, there was a lower average cells/microcarrier for both timepoints in passage 7 compared to passage 4. Additionally, larger aggregates were seen in passage 4 than passage 7 at day 7, and the average cells/microcarrier of aggregates > 50 microcarriers was 5.60 and 4.20 for passage 4 day 7 and passage 7 day 7, respectively. The average single cell volumes quantified by CellTracker Green fluorescence and elastic scattering showed similar overall trends (Figure 4b); there was little change in the average cell volume throughout passage 4, but passage 7 cells were larger in volume overall and actually decreased in volume from day 3 to day 7. The data from off-line study of microcarrier cell growth shows that CellTracker Green and elastic scattering data allow quantification of cell volume. Both fluorescence and elastic scattering modalities showed a linear correlation between total cell volume and nuclear fluorescence-validated cell number (Figure 4c-f). The modalities showed almost equivalent goodness-of-fit values; 0.98 at passage 4 and 0.93 at passage 7.Exclusion of the two outliers (red circles) at passage 7 raises the fluorescence and elastic scattering cell volume R^2^ to 0.9979 and 0.9917, respectively. These outliers could have lower volumes than expected due to their size causing uneven illumination, striping or depth-induced artifacts.

**Figure 4.**
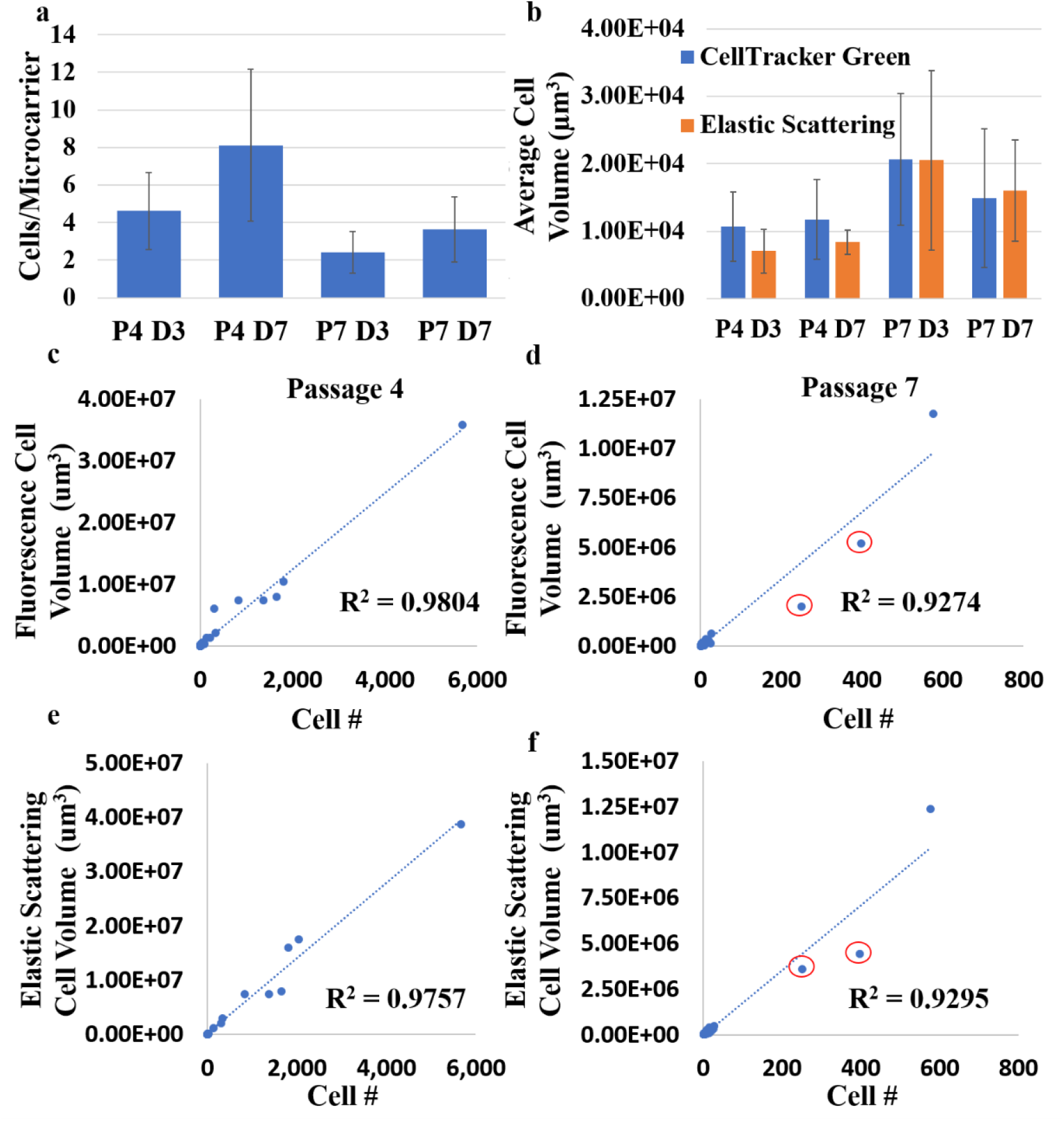
Optical imaging and semi-automated image analysis enable rapid characterization of microcarrier-expanded cell growth *in toto*. **a)** The average CPM at each sampled timepoint for all microcarriers sampled including aggregates from the DRAQ-5 data. **b)** The average single cell volume quantified by the fluorescence-based and the elastic scattering-based methods. The DRAQ-5 labeled cell number versus the total cell volume quantified from the CellTracker Green fluorescence for **c)** passage 4 and **d)** passage 7. The DRAQ-5 labeled cell number versus the total cell volume quantified by the ELIAS method for **e)** passage 4 and **f)** passage 7. The linear trends and R^2^ values are shown. Outliers are identified (red circle).

## Discussion

These experiments first, demonstrate that the off-line fluorescence-based direct cell enumeration and cell volume quantification method can be performed for large microcarrier aggregates, and second, that the ELIAS method can be utilized to non-destructively monitor microcarrier-bioreactor cell culture growth *in toto*, without the need for cellular detachment, membrane lysing, or exogenous fluorescent markers. Due to the refractive index mismatch between the gelMA microcarrier, surrounding agarose, and cell matter, all 3 classes of objects scatter at varying intensities and therefore can be segmented from each other. This allows, for the first time, direct enumeration of cells and microcarriers and quantification of cell volume in a microcarrier culture aggregate using fluorescence and label-free optical imaging modalities (S6 Fig).

Only a handful of studies have focused on the development of optical imaging systems and methods for studying microcarrier-based cell cultures. Jakob et al. used confocal microscopy and LSFM to image MDCK-II cells on Cytodex 3 microcarriers, but only acquired half microcarrier stacks and required sample rotation to visualize cells along the entire microcarrier surface [63]. The optical projection tomography methodology used in Jakob et al. increases the acquisition time and amount of data needed to accurately reconstruct the 3D cell-microcarrier sample compared to more conventional z-stacks for 3D data reconstruction. Duchi et al. demonstrated that optical sectioning via LSFM enables imaging of small cell-microcarrier clumps, but did not image or analyze clumps or aggregates of more than 5 microcarriers as their study focused on single cell motility and distribution on individual microcarriers [74]. *In situ* microscopy and micro-flow imaging enable direct enumeration of cells attached to microcarriers, but both methods are widefield techniques limited to visualizing the proximal half of the microcarrier and neither has the spatial resolution nor contrast for single cell visualization [43], [48]. Odeleye et al., using a custom *in situ* microscope, was only able to image and analyze the proximal microcarrier surface and struggled to enumerate cells accurately once aggregation began [75]. Microflow imaging was used to broadly characterize microcarrier confluency of single microcarriers and small clumps, but the authors did not investigate the ability to analyze large aggregates [76]. Similarly, the map projection analysis method used by Baradez and Marshall to characterize cell growth on individual microcarriers from confocal microscopy data is likely unfeasible for large cell-microcarrier aggregates with true 3D structure [77]. Imaging cytometry presents an attractive solution for on-line visualization and quantification of cells attached to microcarriers, but few studies have used imaging cytometry characterized cells attached to spherical microcarriers or large cell aggregates [46], [47], [78].

The linear relationship between cell number and cell volume quantified by non-destructive *in toto* ELIAS provides confirmation that this method could be used to image and characterize samples in an aqueous environment. An ELIAS PAT for on-line monitoring of microcarrier cell cultures would further incorporate microfluidic chips and hydraulic flow to sample microcarriers from the bioreactor to the lightsheet for rapid analysis and back to the bioreactor culture [79], [80]. Furthermore, single objective lightsheet systems or oblique plane microscopy could better enable on-line imaging of microfluidic samples as there is only a single sample-facing objective and more space for sample mounting and translation below or above the objective [81], [82]. Additional motivation is presented with development of microfluidic chips composed of a polymer with a refractive-index matched to water which is compatible with the presented ELIAS method [83]. These systems and methods are more complex than traditional sampling and imaging methods, but would enable non-destructive, robust, and automated analysis of microcarrier-based and other suspension cultures. Once a cell culture process for cytotherapy manufacturing is standardized and the trend line between off-line fluorescence-validated cell enumeration, on-line ELIAS-quantified, and cytotherapy product quality is confirmed, deviations from the trend line could indicate issues with the culture health and quality. As few as 10-20 populated microcarriers may be needed to quantify the average cells per microcarrier throughout a culture [43].

The *in situ* and *in toto* study of large cell aggregates has, up to this point, been minimal due to the limited imaging depth of conventional microscopes and the opacity of large cell-microcarrier aggregates [75], [77], [84], [85]. Additionally, large aggregates are notoriously difficult to manipulate for study and can cause sampling errors [43], [86]. *In toto* or *in situ* volumetric imaging of these structures would enable analysis of cell morphology, density, and spatial distribution of cell viability throughout an aggregate [30]. Fortunately, the optical sectioning and the decoupled illumination and detection arms of LSM enable imaging deep (>>1 mm) into aggregates. Moreover, the combination of water-dipping objective lens with a long working distance (2 mm), objective lens axial correction collar to fine tune the refractive index mismatch, and hydrogel microcarriers allows for sub-micron resolution volumetric optical imaging of cells attached to spherical microcarriers *in toto*. Because all the pixels in a frame are acquired in parallel, acquisition time can still be relatively short (∼1-2 minutes/4D dataset) even for large aggregates >1 mm in depth. The stripe artifacts, which arise from scattering and/or absorption of the illumination beam by small objects such as air bubbles, debris, or un-melted agarose particles, in LSM images can be removed via a number of hardware and image-processing methods, such as digital scanned lightsheet microscopy (DSLM) with pivot scanning or median digital filtering [87]. Future image analysis pipelines could include corrections for uneven illumination, striping artifacts, or photobleaching [88]. Photobleaching, a chemical alteration of fluorophores that render them unable to fluoresce, is not a concern in esLSM as it does not involve an energy transition; however, extra care has to be taken in esLSM to utilize the full dynamic range of the detector without saturation, even with very low incident laser power.

Standard widefield microscopes, routinely used to visualize and qualitatively evaluate cell culture health, integrate photons from in- and out-of-focus planes, making them suitable for thin samples. Dimensionality reduction, by using a single 2D image for volumetric readouts of 3D cell cultures, leads to cell enumeration and segmentation errors that increase with cell growth and cell-microcarrier aggregates [58], [60], [86]. Cell volume could not be quantified from a single 2D projection of the microcarrier. Volumetric microscopy does enable single cell morphological measurements, such as sphericity or nuclear-to-cytoplasmic volume ratio, for low confluency microcarriers where cells do not overlap (S8 Video). More accurate 3D segmentation methods are needed for single cell segmentation and characterization at moderate or high confluency levels where cells overlap. High-throughput LSM enabling single cell morphological monitoring and profiling could be performed with a more robust cell segmentation method for 3D microcarrier cell cultures [46], [89]. In this study, there was a decrease in cell proliferation and an increase in cell size in passage 7 compared to passage 4, which may be due to replicative senescence [90], [91]. Additional image analysis and more frequent culture sampling could be undertaken to see if time-dependent senescence can be visualized and characterized for microcarrier cultures, as has been investigated for monolayer cultures [32], [33], [92]. Although high-throughput single cell morphological cytometry was not performed here, the presented fluorescence- and elastic-scattering cell growth characterization methods preserve all the spatial (3D cell morphology, distribution) information that is lost to cell lysing and detachment (S9 Video). The ability to characterize single cell morphology will better facilitate real-time decision making regarding the health, quality, or futility of a cell culture process.

## Conclusion

We present optical imaging and image analysis methods for direct cell enumeration and cell volume quantification for microcarrier-expanded cells using *in toto* LSFM and esLSM. Both academic researchers and industry cytotherapy manufacturers would greatly benefit from the ability to monitor and quantify cell culture growth using a non-destructive imaging-based PAT. To the best of our knowledge, this is the first time direct cell enumeration of microcarrier culture aggregates has been reported in the literature.

We used LSM to characterize the entire microcarrier surface, whereas other imaging-based microcarrier growth monitoring methods require either cell membrane lysing or cell-microcarrier detachment, or can only study the proximal half of the microcarrier. Also, we illustrate that our gelMA microcarriers have superior optical imaging capabilities that allow for reliable cell culture monitoring. By incorporating esLSM and hydrogel microcarriers, the ELIAS method presents a strong proof of concept for a non-destructive PAT for monitoring of cytotherapy manufacturing critical parameters. The addition of refractive-index matched microfluidic chips and hydraulic flow would enable on-line ELIAS monitoring of microcarrier-based cell cultures.

## Supporting Information

**S1 Fig. Illustration of the custom 3D-printed sample chamber for imaging on the Zeiss Lightsheet microscope. a)** The chamber allows for dual-sided lightsheet and trans-illumination. **b)** The chamber can be scanned in all 3 dimensions and rotated for stitching and precise sample positioning for 3D optical imaging. Not to scale.

**S2 Fig. Image processing workflow for direct cell enumeration and segmentation of cell nuclei using DRAQ-5 nuclear fluorescence**. The segmented cell nuclei can be analyzed and classified by, for example, average distance to 3 nearest neighbors.

**S3 Fig. Image processing workflow for cell segmentation and quantification using CellTracker Green LSFM data**. Single cells can be classified by volume using Imaris’ morphological-based segmentation.

**S4 Fig. Illustration of image processing workflow for label-free cell segmentation and quantification using elastic scattering LSM data**. Cells scatter at higher intensity values than the microcarrier and surrounding agarose. After the high-pass intensity filter, the voxel filter removes small scatterers or debris in the agarose.

**S5 Fig. Illustration of image processing workflow for microcarrier segmentation and enumeration using elastic scattering LSM data**. The hydrogel microcarriers scatter less than the cells and surrounding agarose. A low-pass intensity and 90 μm size filter identify individual spherical microcarriers. The *en face* and orthogonal cross-sections permit visualization of the gelMA microcarriers using elastic scattering contrast.

**S6 Fig. Resulting multi-modal microscopy data**. Max intensity projections of a passage 7 day 7 microcarrier using the raw **a)** DRAQ-5, **b)** CellTracker Green, and **c)** elastic scattering data. **d)** Merge of CTG + DRAQ-5 fluorescence projections. **e)** Merge of the segmented microcarrier, CTG (shaded green), and DRAQ-5 (shaded blue) cell regions. **f)** Merge of the segmented microcarrier, elastic scattering (shaded white), and DRAQ-5 cell volumes. Scale bar = 25 μm.

**S1 Video. 3D LSFM rendering of a passage 7 day 7 microcarrier with several cells**. No preprocessing was performed on these data. A representative volume to illustrate the data quality of lightsheet microscopy and hydrogel microcarriers (CellTracker Green (green) and DRAQ-5 (blue)). Gamma correction for visualization of the microcarrier only.

**S2 Video. Z-stack scan of passage 7 day 7 microcarrier using LSFM**.

**S3 Video. 3D esLSM rendering of segmented cells attached to the passage 7 day 7 microcarrier**. Elastic scattering contrast permits visualization of cells attached to spherical hydrogel microcarriers similar to that of fluorescence-based contrast.

**S4 Video. Z-stack scan of the esLSM volume illustrating the varying scattering intensities of the cells, microcarriers, and surrounding agarose**. The gelMA microcarrier appear as a solid, non-scattering sphere in the esLSM modality. This allows for direct enumeration of microcarriers.

**S5 Video. 3D fluorescence LSM rendering showing a single cell invading the interior of the hydrogel microcarrier**. CellTracker Green (green) and DiI (gold) show the cell process extending into the center of the microcarrier. DRAQ-5 (blue) apparently resolves two elliptical nuclei within the same nuclear envelope.

**S6 Video. 3D elastic scattering LSM rendering of the same cell invading the core of a microcarrier**. Elastic scattering contrast is able to visualize higher-resolution interesting biological phenomena.

**S7 Video. Z-stack scan of a large microcarrier clump showing CellTracker Green and elastic scattering both enable visualization of complex cell networks that form aggregates**. DRAQ-5-labeled nuclei can be used to semi-automatically large aggregates.

**S8 Video. 3D rendering of passage 7 day 7 segmented microcarrier, CTG cytoplasm, and DRAQ-5 nuclei regions**. No pre-processing performed to segment the cell bodies, nuclei, and microcarrier using CTG, DRAQ-5, and elastic scattering contrast, respectively. These segmented regions are used to quantify cell and nuclear volume, as well as direct cell and microcarrier enumeration.

**S9 Video. 3D rendering of passage 7 day 7 segmented microcarrier, elastic scattering cell body, and DRAQ-5 nuclei regions**. No pre-processing performed to segment the microcarrier, cell bodies, and cell nuclei using elastic scattering and DRAQ-5 contrast. The elastic scattering data enables visualization and detection of microcarriers as well as quantification of cell volume.

## Acknowledgements

We thank the Texas A&M University Microscopy and Imaging Center Core Facility (RRID:SCR_022128) for providing the Zeiss Z1 Lightsheet microscope, Imaris image analysis software, and the technical assistance for their use.

## Author Contributions

**Conceptualization:** Oscar R. Benavides, Holly C. Gibbs, Roland Kaunas, Carl A. Gregory, Kristen C. Maitland

**Funding acquisition:** Holly C. Gibbs, Roland Kaunas, Carl A. Gregory, Kristen C. Maitland

**Investigation:** Oscar R. Benavides, Berkley P. White

**Supervision**: Kristen C. Maitland

**Visualization:** Oscar R. Benavides

**Writing – original draft**: Oscar R. Benavides

**Writing – review and editing:** Oscar R. Benavides, Holly C. Gibbs, Berkley P. White, Roland Kaunas, Carl A. Gregory, Kristen C. Maitland

## References

1. Department of Health and Human Services. PHARMACEUTICAL CGMPS FOR THE 21 ST CENTURY-A RISK-BASED APPROACH FINAL REPORT. Published online 2004. Available from: https://www.fda.gov/media/77391/download

2. Food and Drug Administration. Guidance for Industry PAT - A Framework for Innovative Pharmaceutical Development, manufacturing, and Quality Assurance. Published online 2004. Available from: https://www.fda.gov/media/71012/download

3. Yu LX, Amidon G, Khan MA, Hoag SW, Polli J, Raju GK, et al. Understanding Pharmaceutical Quality by Design. AAPS J. 2014;16(4):771. doi:10.1208/S12248-014-9598-3

4. Lipsitz YY, Timmins NE, Zandstra PW. Quality cell therapy manufacturing by design. Nature Biotechnology 2016 34:4. 2016;34(4):393–400. doi:10.1038/nbt.3525

5. Kirouac DC, Zandstra PW. The systematic production of cells for cell therapies. Cell Stem Cell. 2008;3(4):369–381. doi:10.1016/J.STEM.2008.09.001

6. Boncoraglio GB, Ranieri M, Bersano A, Parati EA, Giovane C del. Stem cell transplantation for ischemic stroke. Cochrane Database Syst Rev. 2019;2019(5). doi:10.1002/14651858.CD007231.PUB3

7. Kondo Y, Toyoda T, Inagaki N, Osafune K. iPSC technology-based regenerative therapy for diabetes. J Diabetes Investig. 2018;9(2):234. doi:10.1111/JDI.12702

8. Antebi B, Pelled G, Gazit D. Stem cell therapy for osteoporosis. Curr Osteoporos Rep. 2014;12(1):41–47. doi:10.1007/S11914-013-0184-X

9. Mohanty R, Chowdhury CR, Arega S, Sen P, Ganguly P, Ganguly N. CAR T cell therapy: A new era for cancer treatment (Review). Oncol Rep. 2019;42(6):2183–2195. doi:10.3892/OR.2019.7335

10. Connon CJ. Bioprocessing for cell based therapies. Wiley Blackwell; 2017. doi: 10.1002/9781118743362

11. Koh B, Sulaiman N, Fauzi MB, Law JX, Ng MH, Idrus RBH, et al. Three dimensional microcarrier system in mesenchymal stem cell culture: A systematic review. Cell Biosci. 2020;10(1):1–16. doi:10.1186/S13578-020-00438-8/TABLES/2

12. McKee C, Chaudhry GR. Advances and challenges in stem cell culture. Colloids Surf B Biointerfaces. 2017;159:62–77. doi:10.1016/J.COLSURFB.2017.07.051

13. Anton D, Burckel H, Josset E, Noel G. Three-Dimensional Cell Culture: A Breakthrough in Vivo. International Journal of Molecular Sciences 2015, Vol 16, Pages 5517-5527. 2015;16(3):5517–5527. doi:10.3390/IJMS16035517

14. Jensen C, Teng Y. Is It Time to Start Transitioning From 2D to 3D Cell Culture? Front Mol Biosci. 2020;7:33. doi:10.3389/FMOLB.2020.00033/BIBTEX

15. Krutty JD, Dias AD, Yun J, Murphy WL, Gopalan P. Synthetic, Chemically Defined Polymer-Coated Microcarriers for the Expansion of Human Mesenchymal Stem Cells. Macromol Biosci. 2019;19(2):1800299. doi:10.1002/MABI.201800299

16. Dwarshuis NJ, Song HW, Patel A, Kotanchek T, Roy K. Functionalized microcarriers improve T cell manufacturing by facilitating migratory memory T cell production and increasing CD4/CD8 ratio. BioRxiv [Prepint]. bioRxiv 646760 [published online May 23, 2019]. Available from: https://www.biorxiv.org/content/10.1101/646760v1.full doi:10.1101/646760

17. Lai JY, Ma DHK. Ocular biocompatibility of gelatin microcarriers functionalized with oxidized hyaluronic acid. Materials Science and Engineering: C. 2017;72:150–159. doi:10.1016/J.MSEC.2016.11.067

18. Vyas KN, Palfreyman JJ, Love DM, Mitrelias T, Barnes CHW. Magnetically labelled gold and epoxy bi-functional microcarriers for suspension based bioassay technologies. Lab Chip. 2012;12(24):5272–5278. doi:10.1039/C2LC41022B

19. Lambrechts T, Papantoniou I, Viazzi S, Bovy T, Schrooten J, Luyten FP, et al. Evaluation of a monitored multiplate bioreactor for large-scale expansion of human periosteum derived stem cells for bone tissue engineering applications. Biochem Eng J. 2016;108:58–68. doi:10.1016/J.BEJ.2015.07.015

20. Egger D, Schwedhelm I, Hansmann J, Kasper C. Hypoxic Three-Dimensional Scaffold-Free Aggregate Cultivation of Mesenchymal Stem Cells in a Stirred Tank Reactor. Bioengineering. 2017;4(2). doi:10.3390/BIOENGINEERING4020047

21. Hassan MNF bin, Yazid MD, Yunus MHM, Chowdhury SR, Lokanathan Y, Idrus RBH, et al. Large-Scale Expansion of Human Mesenchymal Stem Cells. Stem Cells Int. 2020;2020. doi:10.1155/2020/9529465

22. Zhou L, Kong J, Zhuang Y, Chu J, Zhang S, Guo M. Ex vivo expansion of bone marrow mesenchymal stem cells using microcarrier beads in a stirred bioreactor. Biotechnology and Bioprocess Engineering 2013 18:1. 2013;18(1):173–184. doi:10.1007/S12257-012-0512-5

23. Cunha B, Aguiar T, Carvalho SB, Silva MM, Gomes RA, Carrondo MJT, et al. Bioprocess integration for human mesenchymal stem cells: From up to downstream processing scale-up to cell proteome characterization. J Biotechnol. 2017;248:87–98. doi:10.1016/J.JBIOTEC.2017.01.014

24. Rogers RE, Haskell A, White BP, Dalal S, Lopez M, Tahan D, et al. A scalable system for generation of mesenchymal stem cells derived from induced pluripotent cells employing bioreactors and degradable microcarriers. Stem Cells Transl Med. 2021;10(12):1650–1665. doi:10.1002/SCTM.21-0151

25. Lin-Gibson S, Sarkar S, Elliott JT. Summary of the National Institute of Standards and Technology and US Food And Drug Administration cell counting workshop: Sharing practices in cell counting measurements. Cytotherapy. 2018;20(6):785–795. doi:10.1016/J.JCYT.2018.03.031

26. Deskins DL, Bastakoty D, Saraswati S, Shinar A, Holt GE, Young PP. Human Mesenchymal Stromal Cells: Identifying Assays to Predict Potency for Therapeutic Selection. Stem Cells Transl Med. 2013;2(2):151. doi:10.5966/SCTM.2012-0099

27. Tsai AC, Pacak CA. Bioprocessing of Human Mesenchymal Stem Cells: From Planar Culture to Microcarrier-Based Bioreactors. Bioengineering (Basel). 2021;8(7). doi:10.3390/BIOENGINEERING8070096

28. Lin YM, Lee J, Lim JFY, Choolani M, Chan JKY, Reuveny S, et al. Critical attributes of human early mesenchymal stromal cell-laden microcarrier constructs for improved chondrogenic differentiation. Stem Cell Res Ther. 2017;8(1):1–17. doi:10.1186/S13287-017-0538-X

29. Goh TKP, Zhang ZY, Chen AKL, Reuveny S, Choolani M, Chan JKY, et al. Microcarrier Culture for Efficient Expansion and Osteogenic Differentiation of Human Fetal Mesenchymal Stem Cells. Biores Open Access. 2013;2(2):84. doi:10.1089/BIORES.2013.0001

30. Campbell A, Brieva T, Raviv L, Rowley J, Niss K, Brandwein H, et al. Concise Review: Process Development Considerations for Cell Therapy. Stem Cells Transl Med. 2015;4(10):1155. doi:10.5966/SCTM.2014-0294

31. McBeath R, Pirone DM, Nelson CM, Bhadriraju K, Chen CS. Cell shape, cytoskeletal tension, and RhoA regulate stem cell lineage commitment. Dev Cell. 2004;6(4):483–495. doi:10.1016/S1534-5807(04)00075-9

32. Marklein RA, Klinker MW, Drake KA, Polikowsky HG, Lessey-Morillon EC, Bauer SR. Morphological profiling using machine learning reveals emergent subpopulations of interferon-γ-stimulated mesenchymal stromal cells that predict immunosuppression. Cytotherapy. 2019;21(1):17–31. doi:10.1016/J.JCYT.2018.10.008

33. Klinker MW, Marklein RA, lo Surdo JL, Wei CH, Bauer SR. Morphological features of IFN-γ-stimulated mesenchymal stromal cells predict overall immunosuppressive capacity. Proc Natl Acad Sci U S A. 2017;114(13):E2598–E2607. doi:10.1073/PNAS.1617933114

34. Matsuoka F, Takeuchi I, Agata H, Kagami H, Shiono H, Kiyota Y, et al. Morphology-Based Prediction of Osteogenic Differentiation Potential of Human Mesenchymal Stem Cells. PLoS One. 2013;8(2):e55082. doi:10.1371/JOURNAL.PONE.0055082

35. Myers MA. Direct measurement of cell numbers in microtitre plate cultures using the fluorescent dye SYBR green I. J Immunol Methods. 1998;212(1):99–103. doi:10.1016/S0022-1759(98)00011-8

36. Zipper H, Brunner H, Bernhagen J, Vitzthum F. Investigations on DNA intercalation and surface binding by SYBR Green I, its structure determination and methodological implications. Nucleic Acids Res. 2004;32(12):e103. doi:10.1093/NAR/GNH101

37. Vojinović V, Cabral JMS, Fonseca LP. Real-time bioprocess monitoring: Part I: In situ sensors. Sens Actuators B Chem. 2006;114(2):1083–1091. doi:10.1016/J.SNB.2005.07.059

38. Boon M, Luyben KCAM, Heijnen JJ. The use of on-line off-gas analyses and stoichiometry in the bio-oxidation kinetics of sulphide minerals. Hydrometallurgy. 1998;48(1):1–26. doi:10.1016/S0304-386X(97)00074-1

39. Soo B, Lee SC, Lee SY, Chang YK, Chang HN. High cell density fed-batch cultivation of Escherichia coli using exponential feeding combined with pH-stat. doi:10.1007/s00449-003-0347-8

40. Clementschitsch F, Bayer K. Improvement of bioprocess monitoring: development of novel concepts. Microb Cell Fact. 2006;5:19. doi:10.1186/1475-2859-5-19

41. Marose S, Lindemann C, Ulber R, Scheper T. Optical sensor systems for bioprocess monitoring. Trends Biotechnol. 1999;17(1):30–34. doi:10.1016/S0167-7799(98)01247-5

42. Guez JS, Cassar JP, Wartelle F, Dhulster P, Suhr H. Real time in situ microscopy for animal cell-concentration monitoring during high density culture in bioreactor. J Biotechnol. 2004;111(3):335–343. doi:10.1016/J.JBIOTEC.2004.04.028

43. Odeleye AOO, Castillo-Avila S, Boon M, Martin H, Coopman K. Development of an optical system for the non-invasive tracking of stem cell growth on microcarriers. Biotechnol Bioeng. 2017;114(9):2032–2042. doi:10.1002/BIT.26328

44. Hsu CYM, Walsh T, Borys BS, Kallos MS, Rancourt DE. An Integrated Approach toward the Biomanufacturing of Engineered Cell Therapy Products in a Stirred-Suspension Bioreactor. Mol Ther Methods Clin Dev. 2018;9:376–389. doi:10.1016/J.OMTM.2018.04.007

45. He MYC, Stacker SA, Rossi R, Halford MM. Counting nuclei released from microcarrier-based cultures using pro-fluorescent nucleic acid stains and volumetric flow cytometry. Biotechniques. 2017;63(1):34–36. doi:10.2144/000114568/ASSET/IMAGES/LARGE/FUTURE2.JPEG

46. Barteneva NS, Fasler-Kan E, Vorobjev IA. Imaging flow cytometry: coping with heterogeneity in biological systems. J Histochem Cytochem. 2012;60(10):723–733. doi:10.1369/0022155412453052

47. Smith D, Herman C, Razdan S, Abedin MR, Stoecker W van, Barua S. Microparticles for Suspension Culture of Mammalian Cells. ACS Appl Bio Mater. 2019;2(7):2791–2801. doi:10.1021/ACSABM.9B00215/ASSET/IMAGES/LARGE/MT-2019-00215D_0005.JPEG

48. Farrell CJ, Cicalese SM, Davis HB, Dogdas B, Shah T, Culp T, et al. Cell confluency analysis on microcarriers by micro-flow imaging. Cytotechnology. 2016;68(6):2469. doi:10.1007/S10616-016-9967-0

49. Marose S, Lindemann C, Scheper T. Two-dimensional fluorescence spectroscopy: a new tool for on-line bioprocess monitoring. Biotechnol Prog. 1998;14(1):63–74. doi:10.1021/BP970124O

50. Petiot E, Bernard-Moulin P, Magadoux T, Gény C, Pinton H, Marc A. In situ quantification of microcarrier animal cell cultures using near-infrared spectroscopy. Process Biochemistry. 2010;45(11):1832–1836. doi:10.1016/J.PROCBIO.2010.08.010

51. Amini M, Hisdal J, Kalvøy H. Applications of Bioimpedance Measurement Techniques in Tissue Engineering. J Electr Bioimpedance. 2018;9(1):142. doi:10.2478/JOEB-2018-0019

52. Kiviharju K, Salonen K, Moilanen U, Meskanen E, Leisola M, Eerikäinen T. On-line biomass measurements in bioreactor cultivations: comparison study of two on-line probes. J Ind Microbiol Biotechnol. 2007;34(8):561–566. doi:10.1007/S10295-007-0233-5

53. Kiviharju K, Salonen K, Moilanen U, Eerikäinen T. Biomass measurement online: the performance of in situ measurements and software sensors. J Ind Microbiol Biotechnol. 2008;35(7):657–665. doi:10.1007/S10295-008-0346-5

54. Rivera-Ordaz A, Peli V, Manzini P, Barilani M, Lazzari L. Critical Analysis of cGMP Large-Scale Expansion Process in Bioreactors of Human Induced Pluripotent Stem Cells in the Framework of Quality by Design. Biodrugs. 2021;35(6):693. doi:10.1007/S40259-021-00503-9

55. Seiler C, Gazdhar A, Reyes M, Benneker LM, Geiser T, Siebenrock KA, et al. Time-lapse microscopy and classification of 2D human mesenchymal stem cells based on cell shape picks up myogenic from osteogenic and adipogenic differentiation. J Tissue Eng Regen Med. 2014;8(9):737–746. doi:10.1002/TERM.1575

56. Chang WH, Yang ZY, Chong TW, Liu YY, Pan HW, Lin CH. Quantifying cell confluency by plasmonic nanodot arrays to achieve cultivating consistency. ACS Sens. 2019;4(7):1816–1824. doi: 10.1021/acssensors.9b00524

57. Anton F, Burzlaff A, Kasper C, Brückerhoff T, Scheper T. Preliminary study towards the use of in-situ microscopy for the online analysis of microcarrier cultivations. Eng Life Sci. 2007;7(1):91–96. doi:10.1002/ELSC.200620172

58. Bulin AL, Broekgaarden M, Hasan T. Comprehensive high-throughput image analysis for therapeutic efficacy of architecturally complex heterotypic organoids. Sci Rep. 2017;7(1). doi:10.1038/S41598-017-16622-9

59. Salvi M, Morbiducci U, Amadeo F, Santoro R, Angelini F, Chimenti I, et al. Automated Segmentation of Fluorescence Microscopy Images for 3D Cell Detection in human-derived Cardiospheres. Scientific Reports 2019 9:1. 2019;9(1):1–11. doi:10.1038/s41598-019-43137-2

60. Justice C, Leber J, Freimark D, Pino Grace P, Kraume M, Czermak P. Online-and offline-monitoring of stem cell expansion on microcarrier. Cytotechnology. 2011;63(4):325. doi:10.1007/S10616-011-9359-4

61. Bulin AL, Broekgaarden M, Hasan T. Comprehensive high-throughput image analysis for therapeutic efficacy of architecturally complex heterotypic organoids. Sci Rep. 2017;7(1). doi:10.1038/S41598-017-16622-9

62. Celli JP, Rizvi I, Blanden AR, Massodi I, Glidden MD, Pogue BW, et al. An imaging-based platform for high-content, quantitative evaluation of therapeutic response in 3D tumour models. Sci Rep. 2014;4. doi:10.1038/SREP03751

63. Jakob PH, Kehrer J, Flood P, Wiegel C, Haselmann U, Meissner M, et al. A 3-D cell culture system to study epithelia functions using microcarriers. Cytotechnology. 2016;68(5):1813–1825. doi:10.1007/S10616-015-9935-0

64. Huisken J, Swoger J, del Bene F, Wittbrodt J, Stelzer EHK. Optical sectioning deep inside live embryos by selective plane illumination microscopy. Science (1979). 2004;305(5686):1007–1009. doi: 10.1126/science.1100035

65. Nguyen CD, O’Neal PK, Kulkarni N, Yang E, Kang D. Scattering-Based Light-Sheet Microscopy for Rapid Cellular Imaging of Fresh Tissue. Lasers Surg Med. 2021;53(6):872–879. doi:10.1002/LSM.23361

66. Rozbicki E, Downie H, Dupuy LX, MacDonald MP, Yang Z. Light Sheet Tomography (LST) for in situ imaging of plant roots. Optics Express, Vol 21, Issue 14, pp 16239-16247. 2013;21(14):16239–16247. doi:10.1364/OE.21.016239

67. di Battista D, Merino D, Zacharakis G, Loza-Alvarez P, Olarte OE. Enhanced Light Sheet Elastic Scattering Microscopy by Using a Supercontinuum Laser. Methods and Protocols 2019, Vol 2, Page 57. 2019;2(3):57. doi:10.3390/MPS2030057

68. McNeill EP, Zeitouni S, Pan S, Haskell A, Cesarek M, Tahan D, et al. Characterization of a pluripotent stem cell-derived matrix with powerful osteoregenerative capabilities. Nat Commun. 2020;11(1):3025–3025. doi:10.1038/S41467-020-16646-2

69. McNeill EP, Reese RW, Tondon A, Clough BH, Pan S, Froese J, et al. Three-dimensional in vitro modeling of malignant bone disease recapitulates experimentally accessible mechanisms of osteoinhibition. Cell Death & Disease. 2018;9(12):1–18. doi:10.1038/s41419-018-1203-8

70. Benavides OR, Gibbs HC, Gregory CA, Maitland KC. Custom Imaging Chamber for Multimodal Volumetric Microscopy. Biophotonics Congress 2021 (2021), paper DTh2A4. Published online April 12, 2021:DTh2A.4. doi:10.1364/BODA.2021.DTH2A.4

71. Rosenblatt JI, Hokanson JA, Mclaughlin SR, Leary JF. Theoretical Basis for Sampling Statistics Useful for Detecting and Isolating Rare Cells Using Flow Cytometry and Cell Sorting. Cytometry. 1997;27(3):233–8. doi: 10.1002/(sici)1097-0320(19970301)27:3<233::aid-cyto4>3.0.co;2-f

72. di Battista D, Merino D, Zacharakis G, Loza-Alvarez P, Olarte OE. Enhanced Light Sheet Elastic Scattering Microscopy by Using a Supercontinuum Laser. Methods and Protocols 2019, Vol 2, Page 57. 2019;2(3):57. doi:10.3390/MPS2030057

73. Miyazaki K, Yano KI, Saitoh H. A fluorescence method to visualize the nuclear boundary by the lipophilic dye DiI. Biosci Biotechnol Biochem. 2020;84(8):1685–1688. doi:10.1080/09168451.2020.1756737

74. Duchi S, Piccinini F, Pierini M, Bevilacqua A, Torre ML, Lucarelli E, et al. A new holistic 3D non-invasive analysis of cellular distribution and motility on fibroin-alginate microcarriers using light sheet fluorescent microscopy. PLoS One. 2017;12(8):e0183336. doi:10.1371/JOURNAL.PONE.0183336

75. Odeleye AOO, Castillo-Avila S, Boon M, Martin H, Coopman K. Development of an optical system for the non-invasive tracking of stem cell growth on microcarriers. Biotechnol Bioeng. 2017;114(9):2032–2042. doi:10.1002/BIT.26328

76. Farrell CJ, Cicalese SM, Davis HB, Dogdas B, Shah T, Culp T, et al. Cell confluency analysis on microcarriers by micro-flow imaging. Cytotechnology. 2016;68(6):2469–2478. doi:10.1007/S10616-016-9967-0

77. Baradez MO, Marshall D. The Use of Multidimensional Image-Based Analysis to Accurately Monitor Cell Growth in 3D Bioreactor Culture. PLoS One. 2011;6(10):e26104. doi:10.1371/JOURNAL.PONE.0026104

78. Gualda EJ, Pereira H, Martins GG, Gardner R, Moreno N. Three-dimensional imaging flow cytometry through light-sheet fluorescence microscopy. Cytometry Part A. 2017;91(2):144–151. doi:10.1002/CYTO.A.23046

79. Paiè P, Bragheri F, Bassi A, Osellame R. Selective plane illumination microscopy on a chip. Lab Chip. 2016;16(9):1556–1560. doi:10.1039/c6lc00084c

80. Rasmi CK, Padmanabhan S, Shirlekar K, Rajan K, Manjithaya R, Singh V, et al. Integrated light-sheet imaging and flow-based enquiry (iLIFE) system for 3D in-vivo imaging of multicellular organism. Appl Phys Lett. 2017;111(24). doi:10.1063/1.5009782

81. Kumar M, Kishore S, Nasenbeny J, McLean DL, Kozorovitskiy Y. Integrated one-and two-photon scanned oblique plane illumination (SOPi) microscopy for rapid volumetric imaging. Opt Express. 2018;26(10):13027. doi:10.1364/oe.26.013027

82. Bouchard MB, Voleti V, Mendes CS, Lacefield C, Grueber WB, Mann RS, et al. Swept confocally-aligned planar excitation (SCAPE) microscopy for high-speed volumetric imaging of behaving organisms. Nat Photonics. 2015;9(2):113–119. doi:10.1038/nphoton.2014.323

83. Han X, Su Y, White H, O’Neill KM, Morgan NY, Christensen R, et al. A polymer index-matched to water enables diverse applications in fluorescence microscopy. Lab Chip. 2021;21:1549–62. doi:10.1039/d0lc01233e

84. Anton F, Burzlaff A, Kasper C, Brückerhoff T, Scheper T. Preliminary Study towards the Use of In-situ Microscopy for the Online Analysis of Microcarrier Cultivations. Eng Life Sci. 2007;7(1):91–96. doi:10.1002/ELSC.200620172

85. Rudolph G, Lindner P, Gierse A, bluma A, Martinez G, Hitzmann B, et al. Online monitoring of microcarrier based fibroblast cultivations with in situ microscopy. Biotechnol Bioeng. 2008;99(1):136–145. doi:10.1002/BIT.21523

86. Walser M, Leibundgut RM, Pellaux R, Panke S, Held M. Isolation of monoclonal microcarriers colonized by fluorescent E. coli. Cytometry Part A. 2008;73(9):788–798. doi:10.1002/cyto.a.20597

87. Ricci P, Gavryusev V, Müllenbroich C, Turrini L, Vito G de, Silvestri L, et al. Removing striping artifacts in light-sheet fluorescence microscopy: a review. Prog Biophys Mol Biol. 2022;168:52–65. doi:10.1016/J.PBIOMOLBIO.2021.07.003

88. Gibbs HC, Mota SM, Hart NA, Min SW, Vernino AO, Pritchard AL, et al. Navigating the light-sheet image analysis software landscape: concepts for driving cohesion from data acquisition to analysis. Front Cell Dev Biol. 2021;9:2790. doi: 10.3389/fcell.2021.739079

89. Starkuviene V, Pepperkok R. The potential of high-content high-throughput microscopy in drug discovery. Br J Pharmacol. 2007;152(1):62–71. doi:10.1038/sj.bjp.0707346

90. Wagner W, Horn P, Castoldi M, Diehlmann A, Bork S, Saffrich R, et al. Replicative senescence of mesenchymal stem cells: A continuous and organized process. PLoS One. 2008;3(5). doi:10.1371/journal.pone.0002213

91. Zhao Q, Gregory CA, Lee RH, Reger RL, Qin L, Hai B, et al. MSCs derived from iPSCs with a modified protocol are tumor-tropic but have much less potential to promote tumors than bone marrow MSCs. Proc Natl Acad Sci U S A. 2015;112(2):530–535. doi:10.1073/PNAS.1423008112

92. Mota SM, Rogers RE, Haskell AW, McNeill E, Kaunas R, Gregory C, et al. Automated mesenchymal stem cell segmentation and machine learning-based phenotype classification using morphometric and textural analysis. Journal of Medical Imaging. 2021;8(01). doi:10.1117/1.jmi.8.1.014503

