## Supplementary figures and images for "Volumetric Imaging of Human Mesenchymal Stem Cells (hMSCs) for Non-Destructive Quantification of 3D Cell Culture Growth"

### Supplementary Figure 1

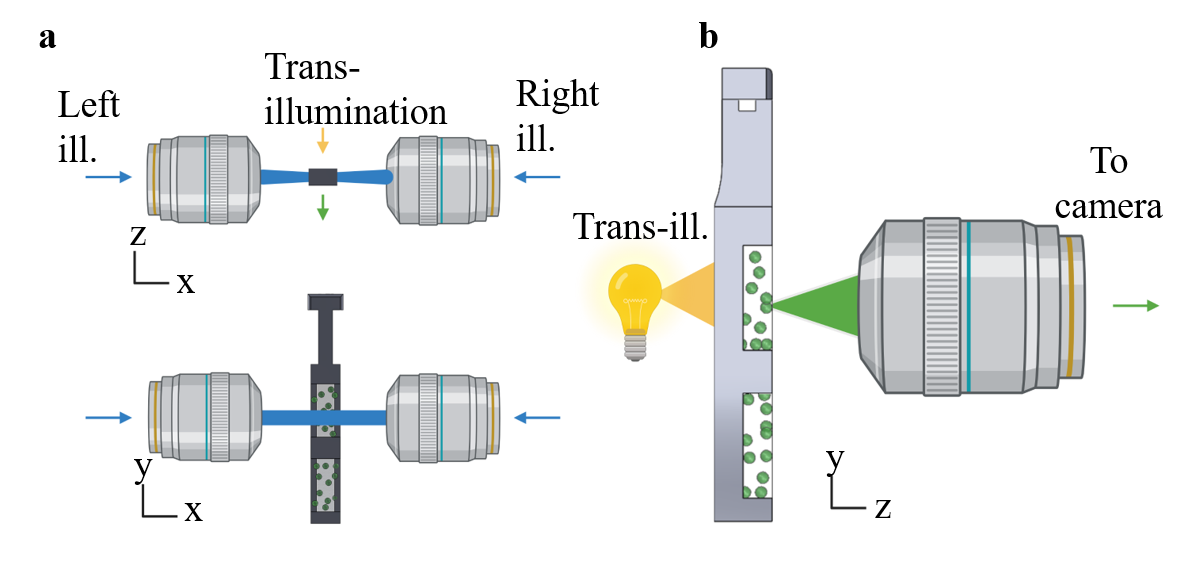

### Supplementary Figure 2

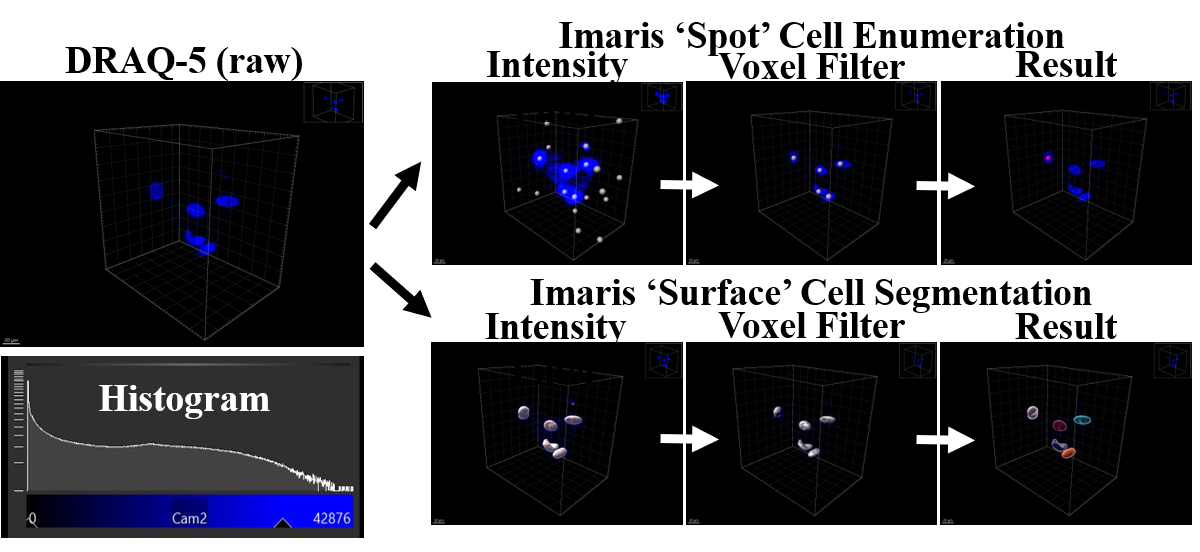

### Supplementary Figure 3

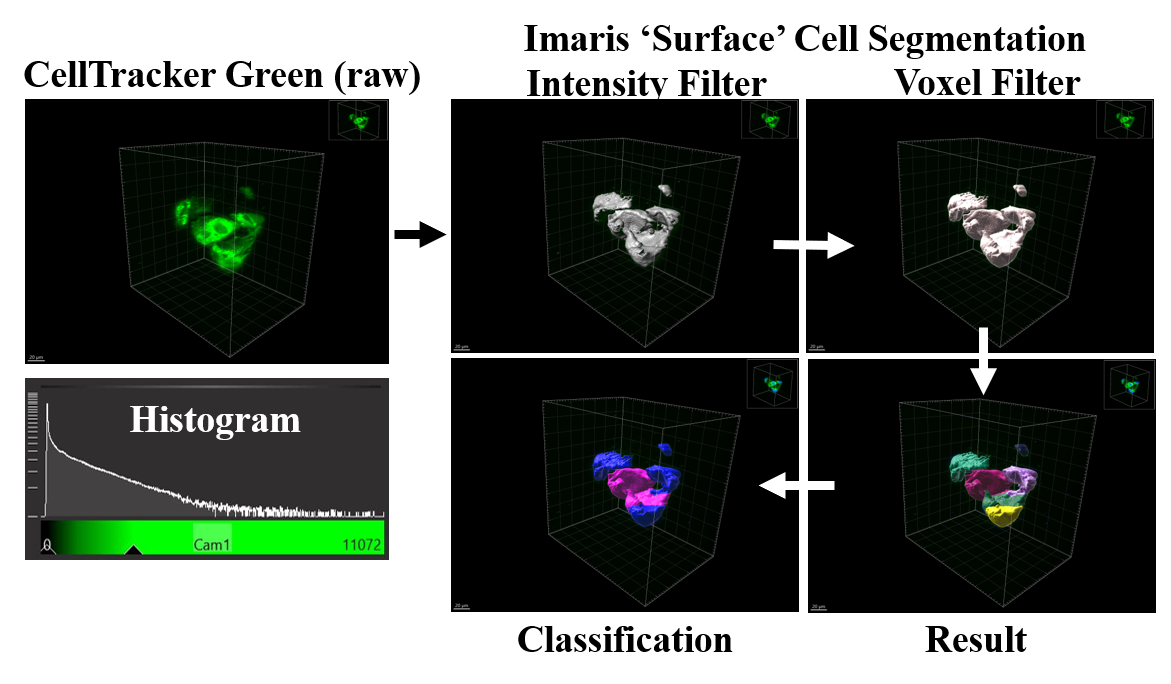

### Supplementary Figure 4

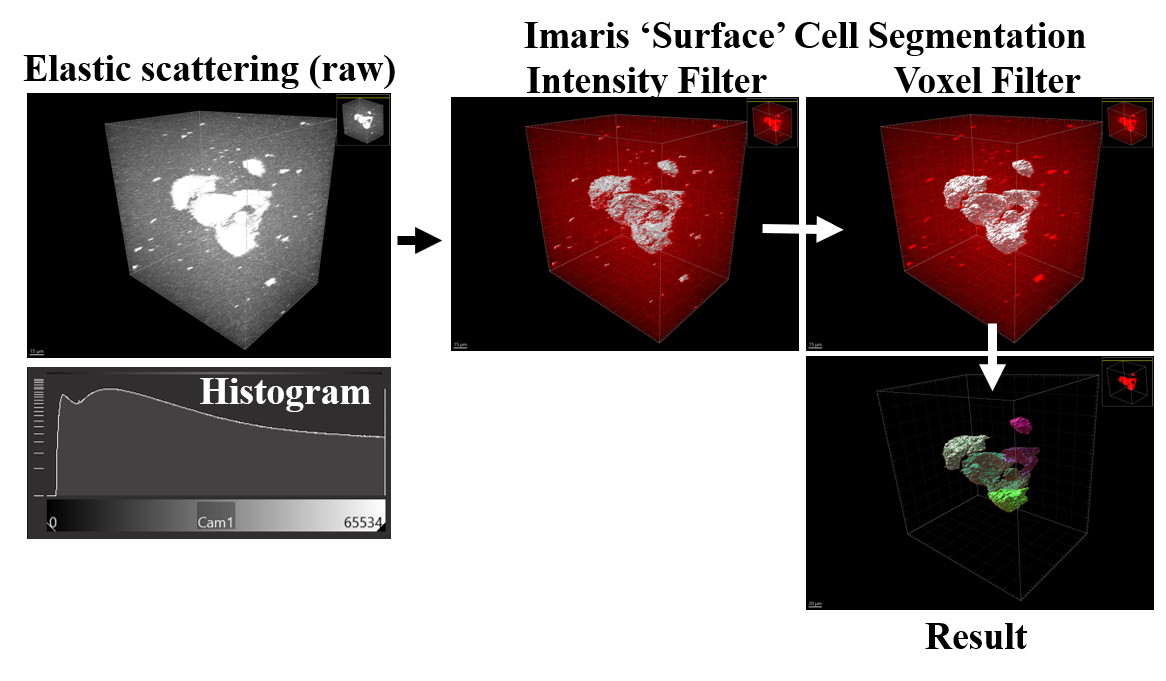

### Supplementary Figure 5

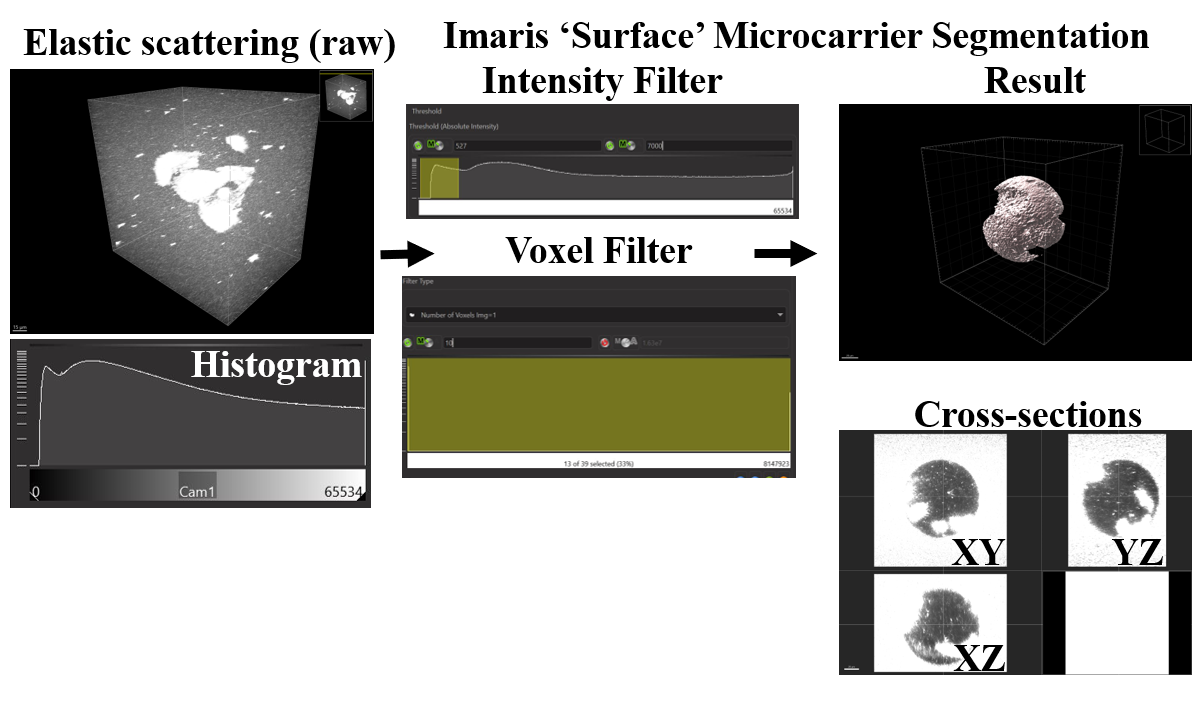

### Supplementary Figure 6

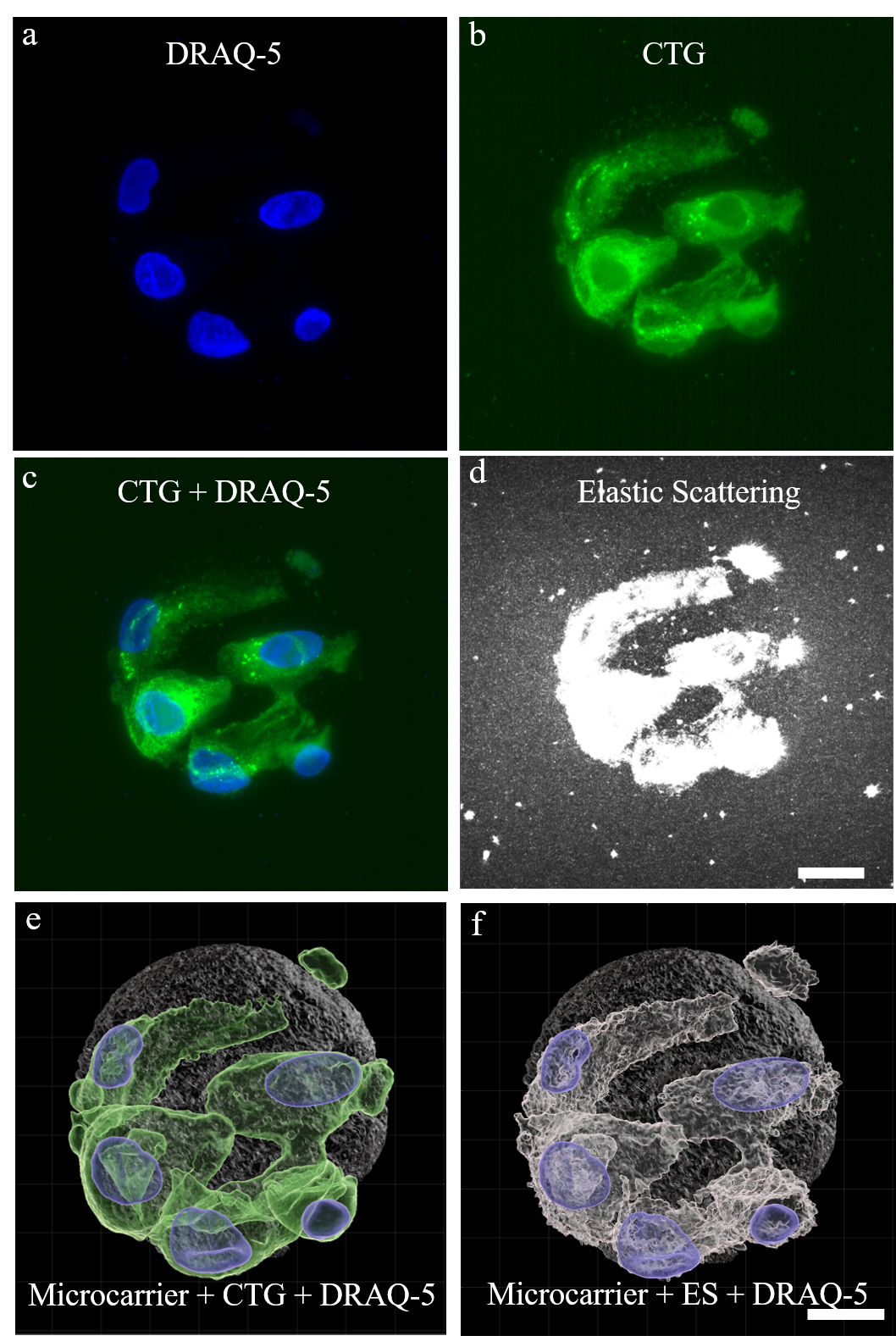
